# VivoFemme: 3D printed human endometrium-on-chip with a thin membrane and gravity-driven perfusion

**DOI:** 10.64898/2026.11.22.768431

**Authors:** Ali Hedayatnia, Arman Jafari, Nour Decae, Ramin Pajoumshariati, Houman Savoji

**Affiliations:** Institute of Biomedical Engineering, Department of Pharmacology and Physiology, Faculty of Medicine, Université de Montréal, Montréal, Quebec H3T 1J4, Canada; Research Center, CHU Sainte-Justine, Montréal, Quebec H3T 1C5, Canada; Montréal TransMedTech Institute (iTMT), Montréal, Quebec H3T 1C5, Canada; BioScience Renal, Research and Early Development, Cardiovascular, Renal and Metabolism, AstraZeneca, Gothenburg 431 83, Sweden; Department of Mechanical Engineering, Polytechnique Montreal, Montréal, Quebec H3T 0A3, Canada; Institute of Biomedical Engineering, Polytechnique Montreal, Montréal, Quebec H3T 0A3, Canada

**Keywords:** endometrium-on-chip, membrane, 3D printing, stereolithography, gravity-driven perfusion, multicellular culture, decidualization, tracer transport

## Abstract

Modeling endometrial function in vitro requires systems that support interactions among epithelial, stromal, and endothelial cells and allow their responses to hormonal and physical cues to be examined. Here, we present VivoFemme, an endometrium-on-chip platform that combines a capillary-pinned gelatin membrane, a stereolithographically 3D-printed microfluidic chip, and gravity-driven perfusion. Gelatin membrane retained on 3D-printed mesh supports separate independently accessible culture micro-channels. The membrane has approximately 10 µm thickness measured by cross-sections using scanning electron microscopy. Optimization of gelatin crosslinking conditions identified a formulation that maintained membrane integrity throughout the culture period. The microfluidic chip operates without external perfusion pumps or circulation tubing and supports primary human endometrial epithelial, stromal, and microvascular endothelial cells in mono-, co-, and tri-culture configurations. Compared with static culture, dynamic endothelial culture showed greater alignment along the flow axis and reduced permeability. Treatment with estradiol, medroxyprogesterone acetate, and cyclic adenosine monophosphate was accompanied by prolactin secretion and insulin-like growth factor-binding protein 1 staining, consistent with stromal decidualization. Prolactin and tracer permeability measurements differed across culture configurations. VivoFemme combines a thin gelatin interface with compartmentalized culture and gravity-driven perfusion to investigate how cellular composition and culture conditions influence endometrial responses.

## 1 Introduction

The human endometrium undergoes changes in epithelial organization, stromal differentiation, and vascular function in response to ovarian hormones. Interactions among these compartments contribute to the establishment of an environment receptive to implantation and supportive of pregnancy^[1]^. Stromal decidualization is central to this process and involves morphological differentiation and changes in secretory activity, including the production of prolactin and insulin-like growth factor-binding protein 1 (IGFBP-1)^[2]^. Single-cell profiling has characterized the diversity and cycle-dependent transitions of endometrial cell populations^[1a, 3]^, while hormone-responsive endometrial organoids have enabled sustained investigation of epithelial biology in vitro^[4]^. Studying interactions among epithelial, stromal, and endothelial cells requires complementary models that bring these populations together under defined culture conditions.

Organ-on-chip technologies allow cells to be organized in spatially defined compartments connected by controlled fluid transport^[5]^. Early epithelial-endothelial models demonstrated that microengineered interfaces could support integrated tissue responses^[6]^, providing a basis for subsequent applications in disease modeling and drug evaluation^[7]^. In reproductive biology, endometrial assembloids combining epithelial organoids and stromal cells have enabled studies of multicellular differentiation and implantation-related processes^[8]^. Multi-organ reproductive-tissue platforms have reproduced a 28-day endocrine profile^[9]^, and dual ovary–endometrium chips have demonstrated bidirectional endocrine communication^[10]^. These approaches provide complementary ways to investigate local cellular interactions and communication between reproductive tissues.

Endometrial microfluidic models have incorporated vascular cells, extracellular matrices, and hormonal stimulation to examine specific aspects of tissue function. Gnecco and colleagues developed a dual-chamber platform with a 6 µm porous resin membrane for stromal–endothelial co-culture^[11]^. Using this platform, they subsequently demonstrated that perfusion of the endothelial compartment enhanced hormone-induced stromal decidualization through paracrine signaling mediated by endothelial-derived prostaglandin E and prostacyclin^[12]^. Ahn and colleagues developed a three-dimensional, fibrin-based endometrium-on-a-chip integrating an epithelial layer, stromal fibroblasts, and self-assembled microvascular networks^[13]^. More recently, this approach was extended to patient-derived epithelial organoids and stromal cells, enabling assessment of endometrial receptivity-associated markers and angiogenic responses^[14]^. De Bem and colleagues used a perfused bovine epithelial–stromal chip to investigate changes in cell-specific transcriptional profiles and secreted protein composition in response to altered glucose and insulin concentrations^[15]^. More recently, Lee and colleagues integrated patient-derived endometrial epithelial organoids and stromal cells with human umbilical vein endothelial cells to assess receptivity-associated molecular markers and angiogenesis^[14]^.

Building on these models, the objective of the present study was to combine a biologically derived interface, directly printed culture compartments, and gravity-driven perfusion in a microfluidic chip that accommodates different endometrial cell types.

Both fabrication and perfusion methods influence how such a device can be constructed and operated. Gravity-driven perfusion supports dynamic culture without external perfusion pumps or circulation tubing^[16]^, while stereolithographic 3D printing enables direct fabrication of microfluidic device geometries^[17]^. Integrating a thin hydrogel interface into a printed microfluidic chip offers a means of separating culture compartments while permitting molecular exchange. Gelatin-derived hydrogels have been used in microengineered cell-culture constructs^[1c, 18]^, motivating the investigation of gelatin-based membrane as interfaces within this architecture.

Matrix composition also influences endometrial cell behavior. Extracellular-matrix biomolecules presented on gelatin-based hydrogels affect endometrial epithelial attachment and cytokeratin expression^[19]^. Synthetic matrices have also supported endometrial organoid–stromal co-culture^[20]^, and hybrid hydrogels containing endometrial extracellular matrix have supported human endometrial organoids^[21]^. Synthetic hydrogels have also enabled microvascularization in an endometriosis microphysiological system^[22]^. These studies emphasize the need to evaluate an interface material within its intended cellular and culture context. The effects of matrix composition, material processing, and device architecture must therefore be considered when interpreting biological responses.

Here, we introduce VivoFemme, an endometrium-on-chip platform that integrates a capillary-pinned gelatin-based membrane, a stereolithographically 3D-printed microfluidic chip, and gravity-driven perfusion. Gelatin membranes retained on 3D-printed mesh supports form interfaces between independently accessible culture micro-channels, with an approximate thickness of 10 µm. The platform accommodates primary human endometrial epithelial, stromal, and microvascular endothelial cells in mono-, co-, and tri-culture configurations, enabling comparisons within a common device architecture.

We characterize membrane formation and stability, swelling and degradation, and the modeled flow conditions, and assess epithelial organization, decidualization-associated markers, tracer transport, and permeability. Compared with static endothelial culture, dynamic culture shows greater alignment along the flow axis and reduced tracer transport and permeability. Hormonal treatment is accompanied by prolactin secretion and IGFBP-1 staining consistent with stromal decidualization, while prolactin measurements and transport responses differ across culture configurations. Together, these experiments assess the application of VivoFemme platform to investigate relationships among cellular composition, flow, and hormone-responsive endometrial behavior.

## 2 Results and Discussion

### Chip architecture and membrane interface

The microfluidic chip was designed in CATIA V5. The microchannel geometry in the microfluidic chip was iteratively evaluated in COMSOL Multiphysics. The device comprises three parallel channels separated by two thin gelatin membranes supported on 3D printed mesh structures (Figure 1A,B). In the tri-culture configuration, stromal and epithelial cells occupy the middle channel, while endometrial microvascular endothelial cells are seeded in the bottom channel. The top channel was not used in this study and could be used for adding another layer (e.g. placenta modeling). Separate access to the channels allows compartment-specific cell seeding and measurements of transport/permeability across the intervening interface.

**Figure 1.**
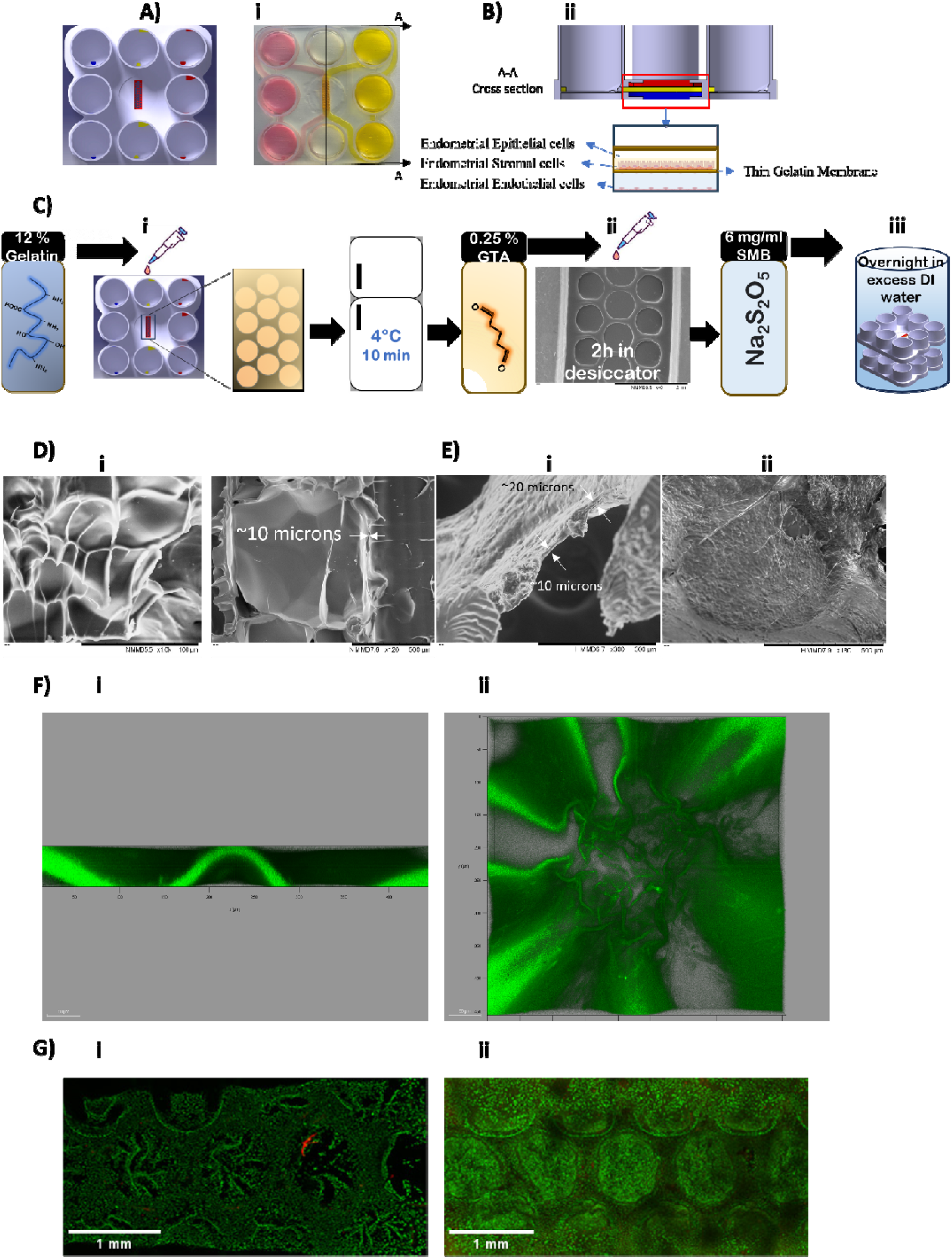
Microfluidic chip architecture and gelatin membrane characterization. A) Microfluidic chip design. B) 3D printed microfluidic chip and schematic cross-section illustrating the three-microchannel configuration. C) Membrane fabrication by capillary pinning of gelatin on the 3D printed mesh, followed by crosslinking and thorough washing. D) SEM images of the gelatin membrane surface and cross-section. E) SEM images showing the membrane with endothelial cell on its surface. F) Confocal images of the hydrated gelatin interface. G) Representative live/dead fluorescence images of primary uterine stromal fibroblasts cultured in the 3D printed microfluidic chip on days 3 and 14. Scale bars are provided in the corresponding image panels.

Capillary pinning retained a gelatin membrane on the mesh after removal of excess precursor solution (Figure 1C). Scanning electron microscopy (SEM) showed a continuous thin membrane, with an approximate thickness of 10 µm in the cross-sections (Figure 1D). Confocal imaging also showed the interface in its hydrated state (Figure 1F), while SEM images showed endothelial cells associated with the membrane surface (Figure 1E).

Postprocessing is an important consideration when preparing microfluidic resin-printed devices for cell culture^[23]^. Following isopropanol washing, post-curing, and MiliQ water rinsing, the microfluidic chips supported primary uterine stromal fibroblast culture, with viable cells visible in live/dead images on days 3 and 14 (Figure 1G). These observations support the biocompatibility of the processed device with stromal-cell culture under the tested conditions.

### Flow modeling and flow-induced endothelial response

Gravity-driven rocking enabled dynamic culture without an external perfusion pump or circulation tubing. Flow was estimated using a draining-tank approximation and compared with measurements obtained from a prototype device. The resulting flow conditions were used as inputs for steady-state, laminar-flow computational fluid dynamics simulations using COMSOL Multiphysics. The wall shear stress (WSS) values in the top and bottom channels were approximately 3-8 dyn cm ² across the modeled conditions (Figure 2A,B). The spatial distributions followed similar patterns at the different flow rates.

**Figure 2.**
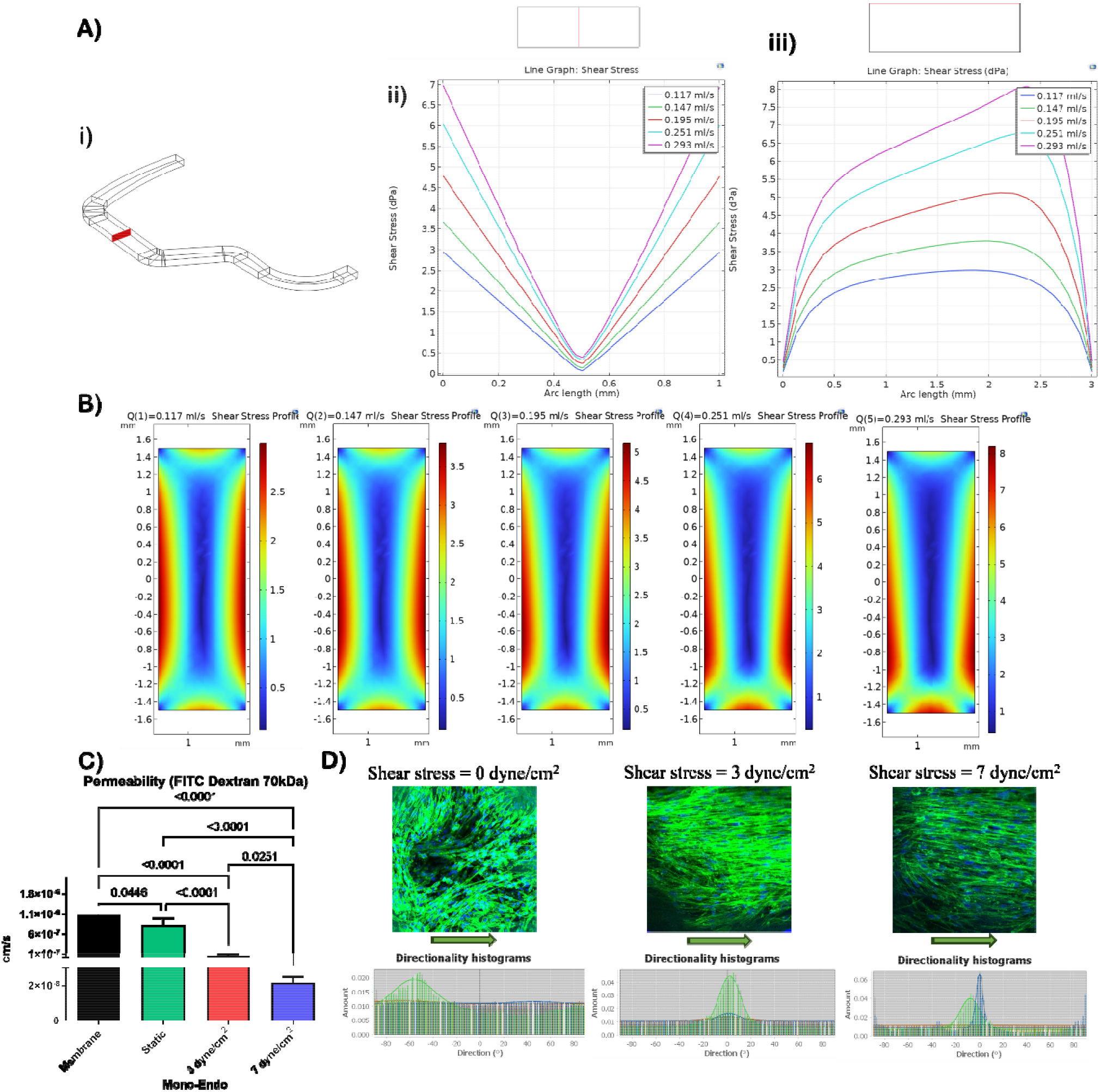
Computational flow characterization and endothelial responses under static and dynamic culture conditions. A) Computational geometry and wall shear stress (WSS) profiles at the modeled flow rates. B) Spatial WSS distributions for the corresponding flow conditions. C) Appa ent permeability to 70 kDa FITC-dextran across a cell-free membrane and endothelial cultures maintained under static or dynamic conditions with nominal WSS values of 3 and 7 dyn cm□*². Data are presented as mean ± SD; displayed values indicate p-values. D) Representative endothelial-cell images and corresponding orientation distributions under the indicated culture conditions. Arrows indicate the flow axis. WSS expressed in dPa is numerically equivalent to dyn cm ². Endo represents endothelial cells*.

Compared with static culture, endothelial cultures maintained under dynamic conditions showed greater alignment along the flow axis and lower apparent tracer (i.e., 70 kDa FITC-dextran) permeability. Apparent permeability decreased further at the higher nominal WSS condition (Figure 2C,D). These observations are consistent with reports of flow-dependent changes in endothelial morphology and barrier properties in endometrial and other microfluidic models^[11, 24]^. In the present device, however, we acknowledge that flow reversal and changes in reservoir levels (inlet and outlet wells) during rocking produce time-dependent conditions are not addressed by the steady-state simulations. Medium replacement can also alter the reservoir levels and, consequently, the flow conditions.

### Membrane fabrication and characterization

Membranes were fabricated from a 12 wt% type B bovine gelatin solution crosslinked with glutaraldehyde. Preliminary optimization identified this gelatin concentration as suitable for retaining films on the 3D printed mesh (data not shown). Membrane stability varied with glutaraldehyde concentration, for example, membranes prepared with 0.125% glutaraldehyde typically ruptured within 2-4 days of cell culture, whereas those prepared with 1.25% glutaraldehyde were brittle during handling. Membranes prepared with 0.25% glutaraldehyde remained intact during the co-culture experiments. Therefore, 0.25% glutaraldehyde concentration was selected for subsequent studies.

Bulk gelatin specimens exhibited approximately 500-800% swelling after 24 h in phosphate-buffered saline (PBS), with lower water uptake at higher glutaraldehyde concentrations (Figure 3B). Specimens prepared with 0.25% glutaraldehyde showed approximately 12% and 16% mass loss after 2 and 4 weeks, respectively (Figure 3A). These observations are consistent with the influence of glutaraldehyde crosslinking on gelatin hydration and stability^[25]^.

**Figure 3.**
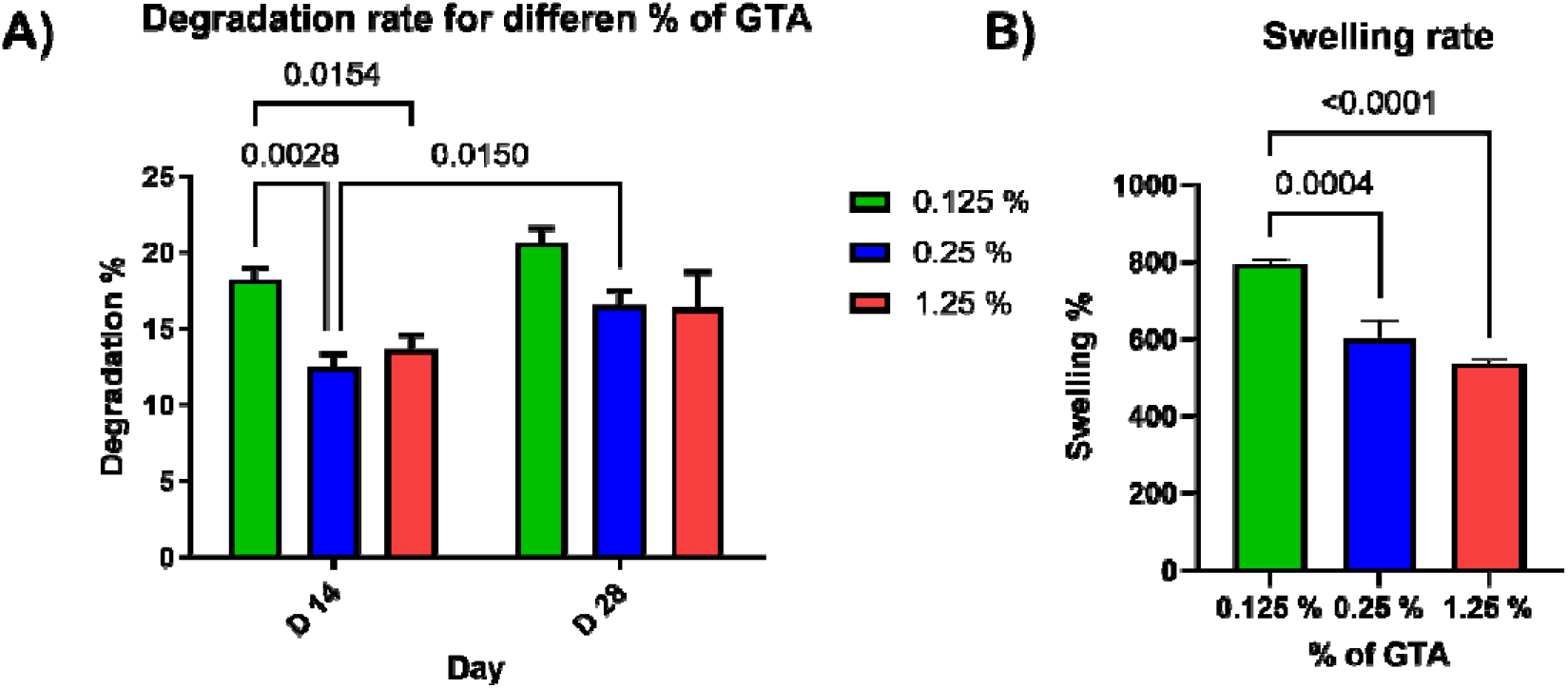
Membrane characterization and epithelial organization under different culture conditions. A) Mass loss of gelatin membranes after 2 and 4 weeks for the indicated glutaraldehyde concentrations. B) Swelling of the corresponding formulations after 24 h in PBS.

Epithelial organization was examined using 10%, 20%, and 50% Geltrex in mono-, co-, and tri-culture configurations. Epithelial monoculture with 20% Geltrex was identified in the initial optimization as the condition yielding the greatest number and size of gland-like structures among the groups tested (data not shown). Based on this optimization, 20% Geltrex was selected for epithelial encapsulation in subsequent experiments.

### Hormonal response and decidualization-associated markers

Cultures were treated with 10 nM estradiol, 1 µM medroxyprogesterone acetate (MPA), and 0.5 mM cyclic adenosine monophosphate (cAMP) from day 1 through day 10. Samples for prolactin measurement were collected on days 3, 5, and 10 (Figure 4A). This regimen draws on established previously published protocols for progestin- and cAMP-responsive endometrial culture^[26]^. Because different decidualization stimuli can produce distinct transcriptional responses in human stromal cells, treatment composition and dose should be considered when comparing outcomes across models^[27]^. Measured prolactin values were lower in on-chip tri-culture than in 2D stromal monoculture (Figure 4B). In the comparison among culture configurations, epithelial-stromal co-culture showed lower prolactin values than stromal monoculture, whereas differences between endothelial-stromal co-culture and stromal monoculture were less pronounced (Figure 4C).

**Figure 4.**
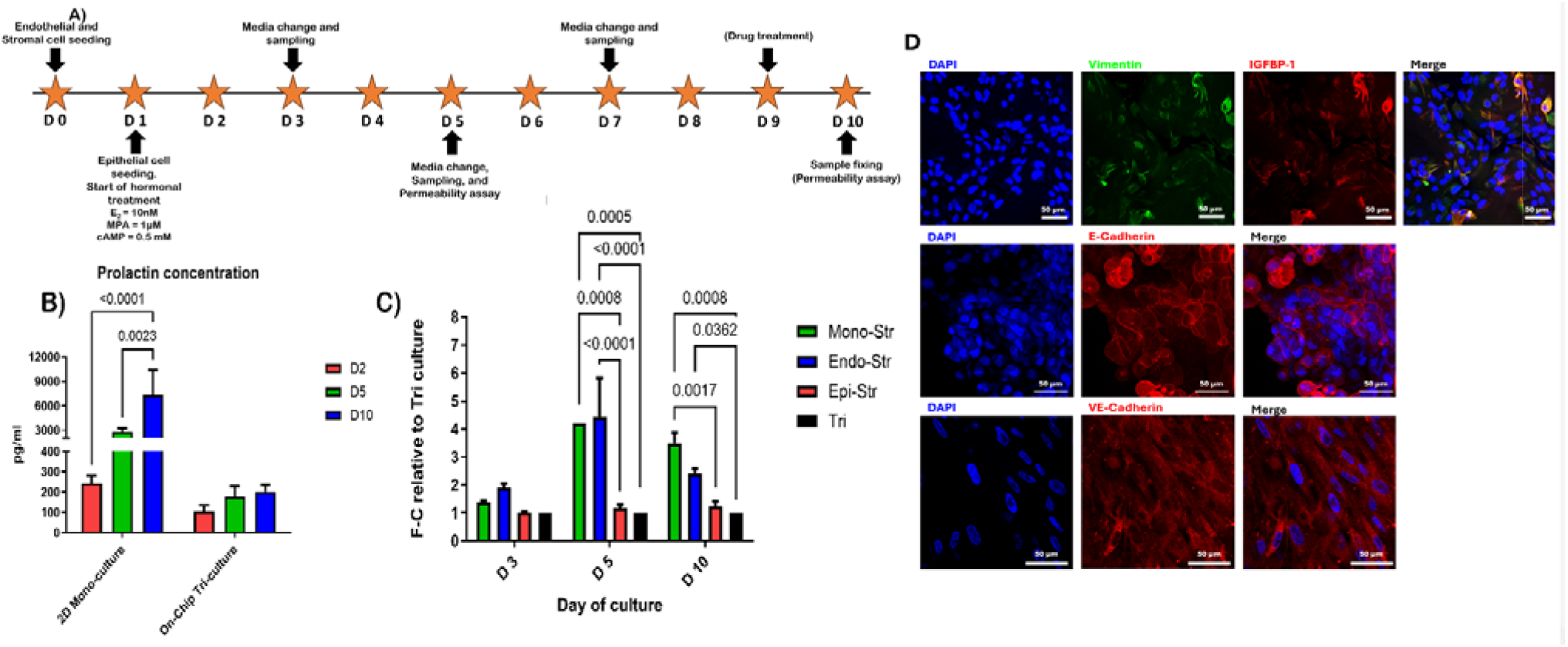
Hormonal treatment, prolactin measurements, and endometrial marker expression. A) *Timeline of cell seeding, treatment, sampling, and imaging. Cultures received 10 nM estradiol, 1 µM medroxyprogesterone acetate (MPA), and 0.5 mM cyclic adenosine monophosphate (cAMP) from day 1 through day 10. B) Prolactin measurements in 2D stromal monoculture and on-chip tri-culture on days 3, 5, and 10. Comparisons used the reported culture-area normalization. C) Prolactin values across culture configurations expressed as fold changes relative to tri-culture. D) Representative fluorescence images after 10 days of culture showing vimentin and insulin-like growth factor-binding protein 1 (IGFBP-1; top row), E-cadherin (middle row), and VE-cadherin (bottom row), with nuclear counterstaining. Scale bars: 50 µm. Mono-Str, stromal monoculture; Endo-Str, endothelial–stromal co-culture; Epi-Str, epithelial–stromal co-culture; Tri, epithelial–stromal–endothelial tri-culture*.

Prolactin comparisons used an area-based normalization, with culture surface areas of 1.9 cm² for the 2D 24-well culture and 1.09 cm² for the stromal-containing chip compartment with equal cell density at confluence. The lower measurements in epithelial-containing cultures may be explained by epithelial-stromal signaling which is in agreement with previous work with endometrial organoid-stromal co-culture in a defined synthetic matrix^[20]^. Moreover, the differences between endothelial-stromal co-culture and stromal monoculture should likewise be interpreted within the flow, medium, and stimulation conditions used here. In a related study, Gnecco and colleagues demonstrated enhanced decidualization through endothelial prostaglandin signaling under continuous perfusion^[12]^, indicating that endothelial contributions can depend on the experimental setting.

IGFBP-1 staining after 10 days of culture provided an additional observation consistent with stromal decidualization (Figure 4D, top row). E-cadherin and VE-cadherin staining showed epithelial and endothelial organization, respectively, at the end of the culture period (Figure 4D, middle and bot om rows). Together with the prolactin measurements, these findings demonstrates the presence of decidualization-associated responses in a multicellular endometrial culture.

### Tracer transport/permeability and pharmacological perturbation

Transport/permeability was assessed using fluorescent inulin, 70 kDa dextran, and 2-NBD-glucose (2-NBDG). In endothelial monoculture, apparent permeability to 70 kDa dextran was lower under dynamic conditions than under static conditions and decreased further at the higher nominal WSS condition (Figure 2C). In tri-culture, the reported analysis detected no statistically significant differences between hormone-treated and untreated constructs for the probes tested (Figure 5A).

**Figure 5.**
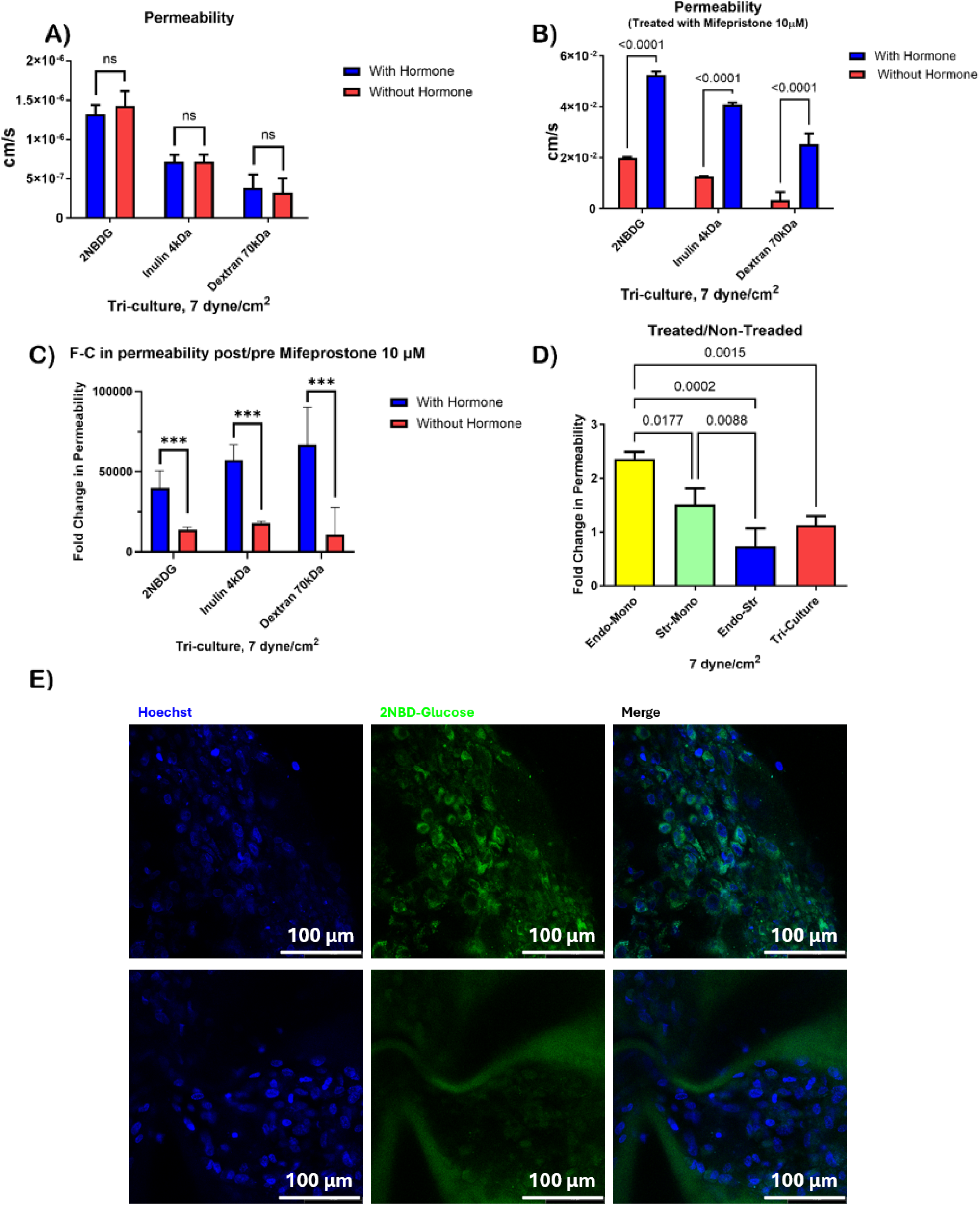
Tracer transport and responses to mifepristone. A) Apparent permeability to 2-NBDG, *fluorescent inulin, and 70 kDa dextran in tri-cultures with and without hormonal stimulation. B) Apparent permeability on day 10 after exposure to 10 µM mifepristone for 24 h and removal of drug-containing medium before the tracer assay. C) Reported fold changes in apparent permeability following mifepristone exposure. D) Ratios of apparent permeability in hormone-treated versus untreated cultures across the indicated cellular configurations. E) Representative fluorescence images showing cell-associated 2-NBDG signal and nuclear counterstaining. Scale bars: 100 µm. Endo, endothelial cells; Str, stromal cells; Endo–Str, endothelial–stromal co-culture; Tri, epithelial–stromal–endothelial tri-culture*.

Across culture configurations, hormone-treated/untreated apparent-permeability ratios were approximately 2.3 in endothelial monoculture and 1.5 in stromal monoculture, whereas the ratios for endothelial-stromal co-culture and tri-culture were closer to 1 (Figure 5D). These differences may reflect cell coverage, matrix resistance to transport, or interactions among cell populations. Due to the combined treatment with estradiol, MPA, and cAMP, the observed responses may be attributed specifically to progesterone receptor signaling. Previous works reported that endothelial progesterone receptor signaling has been linked to physiological preimplantation permeability in the mouse uterus^[28]^. Similarly, decidualization-associated changes in stromal-cell morphology and mechanical properties may influence transport/permeability^[29]^. The measurements in this work therefore describe transport/permeability across the cell-matrix construct.

Representative images showed greater cell-associated 2-NBDG fluorescence in hormone-treated cultures (Figure 5E). This observation is consistent with increased accumulation. In other cell systems, 2-NBDG binding and uptake have been shown to occur independently of membrane glucose transporters^[30]^.

Following exposure to 10 µM mifepristone for 24 h, both hormone-treated and untreated constructs showed large apparent increases in transport/permeability (Figure 5B,C). Measurements were obtained from the same chips on day 5 and again on day 10.

## 3 Conclusion

We successfully developed and evaluated a microfluidic chip designed to replicate the physiological conditions of endometrium. The microfluidic chip design, characterized by microchannel optimization and use of thin ECM membranes, demonstrated significant potential in providing a robust and physiologically relevant environment for cell co-culture. The three-channel design, with ultra-thin ECM membranes, effectively facilitated cell adhesion and growth, closely mimicking physiological conditions.

Integrating CATIA V5 and COMSOL Multiphysics allowed us to create a microchannel geometry that ensures laminar and uniform flow, essential for maintaining endothelial and epithelial cell functions. Flow rate estimations and subsequent CFD simulations revealed that the microchannels’ wall shear stress (WSS) ranged from 3 to 8 dyne/cm², aligning with physiological conditions. 12% gelatin solution crosslinked with 0.25% glutaraldehyde provided an optimal balance of mechanical properties and low cytotoxicity, essential for long-term cell culture. Higher glutaraldehyde concentrations resulted in too stiff membranes, while lower concentrations led to membrane rupture due to cell migration and junction formation. The gelatin hydrogel’s mechanical strength, chemical stability, and active cell binding sites facilitated cell culture without secondary coatings, reducing its application.

Hormonal treatments using estradiol, medroxyprogesterone acetate, and cAMP in tri-culture setups revealed significantly reduced prolactin expression compared to 2D monoculture of stromal cells, highlighting the complex cell interactions in this microenvironment, underscoring the relevance of replicating the complex cellular interactions and hormonal responses of the female reproductive system. Permeability assays demonstrated that endothelial cells cultured under dynamic conditions exhibited reduced permeability, which decreased further with increased WSS. Hormonal treatments increased permeability in endothelial cell monocultures, consistent with known physiological responses to progesterone. The lack of significant permeability changes in co-culture and tri-culture setups highlighted the complex interplay of maintaining barrier integrity under hormonal influence.

The microfluidic chip developed in this study demonstrates a robust and physiologically relevant platform for studying endometrium and in near term female reproductive physiology. The chip’s design ensures appropriate mechanical and biochemical environments for cell multi-culture, while its ability to replicate complex cell interactions and hormonal responses offers valuable insights into reproductive health. These findings highlight the potential of this microfluidic chip in advancing reproductive research.

## 4 Experimental Section

### Device design and fabrication

The device was designed in CATIA V5, and its geometry was evaluated in COMSOL Multiphysics to refine the connecting channels and co-culture region. The dimensions of micro-channels within the co-culture region were 3 mm in width, 1 mm in height, and 12 mm in length. The reported mesh feature diameter was 0.9 mm. The effective membrane exchange area used for transport calculations was 19.72 mm² (0.1972 cm²).

Flow was modeled using a steady-state laminar-flow study with a defined inlet flow rate, a pressure outlet, and a physics-controlled mesh. Flow estimates obtained from a draining-tank model were compared with measurements from a prototype device.

Microfluidic chip devices were fabricated using a Formlabs 3B stereolithography printer and BioMed Clear resin. Printed parts underwent two washes in ≥99% isopropanol, each lasting 1 h, in a Form Wash system, followed by post-curing for 90 min at 60 °C in a Form Cure system. The devices were then immersed overnight in ultrapure water at 60 °C and dried. The co-culture region was sealed with two glass coverslips using Eukitt quick-hardening mounting medium (Sigma). The mounting medium was allowed to cure at room temperature under a fume hood for at least 2 h.

### Membrane preparation and characterization

4 mL of a 12 wt% type B bovine-skin gelatin solution (Sigma-Aldrich, USA) was cast in an 8 cm-diameter Petri dish to produce a gel with a thickness of approximately 500 µm. Samples were held at 4 °C for 15 min before adding 12 mL of glutaraldehyde solution at a nominal concentration of 0.125%, 0.25%, or 1.25%, prepared from a 25% stock. Samples were maintained under vacuum for 20 h. The crosslinker solution was then replaced with 15 mL of sodium metabisulfite solution at 8 mg mL ¹ for a further 20 h, followed by washing.

For swelling measurements, dried samples were immersed in phosphate-buffered saline (PBS) at 37 °C for 24 h, and weighed. Swelling was calculated as:

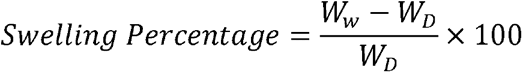

where W is the swollen sample mass and W_D_ is the initial dry sample mass.

For mass-loss measurements, samples were incubated in 12 mL PBS at 37 °C, with PBS replacement every other day. Samples were collected after 14 and 28 days, rinsed with distilled water, dried in an oven at 50 °C, and weighed. Mass loss was expressed as:

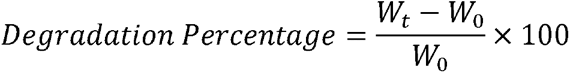

where W□is the initial dry sample mass and W_t_ is the dry sample mass after incubation for time t. Swelling and mass-loss experiments were performed in technical triplicate.

### On-chip membrane formation

Type B bovine-skin gelatin (Sigma-Aldrich, USA) was dissolved in water at 12 wt% by stirring at 60 °C until the solution was homogeneous and free of visible bubbles. Prewarmed chips were allowed to cool to approximately 40 °C before 200 µL of gelatin solution was introduced into the middle channel. Excess solution was gently withdrawn through a pipette tip connected to a vacuum source, leaving a gelatin film pinned to the mesh. The loading and withdrawal procedure was repeated twice.

Chips were held at room temperature for 2 min and then at 4 °C for 10 min. The films were crosslinked for 2 h in a desiccator using 2 mL of nominal 0.25% glutaraldehyde solution prepared from a 25% stock. The crosslinking reaction was quenched with 6 mg mL□¹ sodium metabisulfite for 1 h, followed by overnight extraction in water.

Membrane morphology was examined by scanning electron microscopy (SEM), and the hydrated interface was imaged by confocal microscopy.

### Cell sources and expansion

Human uterine stromal fibroblasts (ScienCell, catalog no. 7040) and human endometrial microvascular endothelial cells (ScienCell, catalog no. 7010) were expanded in fibroblast medium (catalog no. 2301) and endothelial medium (catalog no. 1001), respectively, with supplier-recommended supplements. Human endometrial epithelial cells (Lifeline Cell Technology, catalog no. FC-0078) were expanded in ReproLife reproductive medium (catalog no. LL-0068). Cells were dissociated at approximately 85–100% confluence and used for experiments at passages 3-6.

### Cell seeding and culture configuration

Before cell seeding, chips were incubated for 30 min in PBS containing 2% penicillin–streptomycin. Residual liquid was removed by vacuum suction, and chips were air-dried for 10 min in a biological safety cabinet.

For stromal-cell seeding, a stock suspension containing 2 × 10□cells mL□¹ was mixed with sterile-filtered fibrinogen stock at 10 mg mL□¹ in fibroblast medium at a cell-suspension-to-fibrinogen-stock volume ratio of 0.6:1. This ratio corresponds to nominal concentrations of 7.5 × 10□cells mL□¹ and 6.25 mg mL□¹ fibrinogen before any subsequent dilution or gelation. Thrombin solution (150 µL, 5 U mL ¹) was introduced into the bottom channel for 5 min. The fibrinogen-cell mixture (150 µL) was then introduced into the middle channel and incubated for 10-15 min. Residual precursor and thrombin solutions were gently withdrawn, leaving stromal cells embedded in the fibrin matrix. Stromal monocultures received 2 mL of fresh fibroblast medium. Cultures designated for subsequent seeding with additional cell types initially received 150 µL of medium.

For endothelial seeding, 150 µL of fibronectin solution (0.01 mg mL□¹ in endothelial medium; ScienCell, catalog no. 8248) was introduced into the bottom channel. Chips were inverted and incubated for 30 min to coat the membrane-facing surface. An endothelial-cell suspension at 2.5 × 10 cells mL□¹ was then introduced into the bottom channel at a volume of 150 µL. Chips were immediately inverted and incubated undisturbed for 1 h. The channel was gently perfused by gravity-driven medium flow and supplied with 2 mL of fresh endothelial medium. A fibronectin-coated 24-well plate receiving the same volume of cell suspension served as an attachment control.

For epithelial seeding, 150 µL of 5% Geltrex (Gibco, catalog no. A1413302) in reproductive medium was introduced into the middle channel and incubated at 37 °C for 1 h. Epithelial cells (3 × 10 cells mL□¹) were encapsulated in 20% Geltrex, and 150 µL of the cell-containing suspension was introduced into the channel and incubated for 1 h. An additional 150 µL of 5% Geltrex was applied over the epithelial culture. Chips were left undisturbed for 2-3 h, gently washed by gravity-driven flow, and supplied with 2 mL of reproductive medium before dynamic culture. A 24-well plate receiving the same volume of cell suspension served as an attachment control. Medium was replaced every two days. Screening of gland-like epithelial organization also included 10%, 20%, and 50% Geltrex in the indicated culture configurations (data not shown).

Dynamic cultures were maintained on a CelXrocker operated in tide mode at a reported speed setting of 6 rpm and a tilt setting of 20°, with a reservoir volume of 2 mL.

### Hormonal stimulation

Cultures were treated with 10 nM 17β-estradiol, 1 µM medroxyprogesterone acetate (MPA), and 0.5 mM cyclic adenosine monophosphate (cAMP) from day 1 through day 10. Samples for prolactin measurement were collected on days 3, 5, and 10.

A 100 µM estradiol stock solution was prepared in ethanol and diluted in phenol-red-free DMEM/F12 containing 10% charcoal-stripped fetal bovine serum. A 100 mM MPA stock solution was prepared in dimethyl sulfoxide (DMSO) and diluted in reproductive medium to 1 µM. A 23.251 mM cAMP stock solution was prepared in reproductive medium and diluted to 0.5 mM.

### Transport/permeability assay and mifepristone exposure

Tracer solutions were prepared in reproductive medium: 70 kDa FITC-dextran (Millipore Sigma) and 70 kDa TRITC-dextran (Chondrex, catalog no. 4014) at 1 mg mL□¹; FITC-inulin (approximately 4.5 kDa; catalog identifier FI-100 MG) at 100 µM; and 2-NBDG (Abcam) at 0.1 mM. Permeability was assessed on day 5 and again on the same chips on day 10, after mifepristone exposure and removal.

Before each assay, the channels were rinsed once with PBS. The middle receiver compartment received 2 mL of reproductive medium, and the bottom donor compartment received 2 mL of tracer solution. Chips were maintained on the rocking platform for 2 h. Samples of 100 µL were then collected from the donor and receiver reservoirs and transferred to a black 96-well plate. Tracer concentrations were determined using serial-dilution fluorescence calibration curves prepared for each probe.

For pharmacological perturbation, mifepristone was dissolved in dimethyl sulfoxide (DMSO) and added to culture medium at a final concentration of 10 µM. Hormone-treated and untreated constructs were exposed from day 9 to day 10 for 24 h. Drug-containing medium was removed before the day-10 tracer assay. The day-5 and day-10 measurements therefore constituted longitudinal observations from the same chips.

### Viability and fluorescence imaging

Cell viability was assessed on days 1, 3, and 14 using the reported LIVE/DEAD Viability/Cytotoxicity Kit for mammalian cells (Invitrogen, catalog no. L3224), with calcein AM and a dead-cell counterstain. Staining solution (1 mL) was added to each reservoir, and samples were incubated at 37 °C for 20 min before confocal imaging.

For immunofluorescence, cells were washed three times with PBS, fixed with 4% paraformaldehyde (Thermo Scientific, catalog no. AAJ19943K2) for 30 min at room temperature, and washed three additional times with PBS. Samples were permeabilized with 0.1% Triton X-100 and blocked with 10% goat serum. Primary antibodies were diluted in PBS containing 10% goat serum and incubated with samples overnight at 4 °C. The antibodies were directed against VE-cadherin (D87F2; Cell Signaling Technology, catalog no. 2500; 1:400), E-cadherin (4A2; catalog no. 14472; 1:100), IGFBP-1 (Abcam, catalog no. ab228741; 1:200), and vimentin (D21H3; Cell Signaling Technology, catalog no. 9854; 1:200).

After three PBS washes, samples were incubated with secondary antibodies diluted 1:1000 in PBS containing 1% goat serum for 1 h at room temperature. The reported secondary antibodies were Alexa Fluor 594-conjugated goat anti-rabbit antibody (Abcam, catalog no. ab150080) and Alexa Fluor 647-conjugated anti-mouse antibody (Cell Signaling Technology, catalog no. 4410). Nuclei were counterstained with DAPI, followed by three final PBS washes. Fluorescence images were acquired using a Leica SP8 inverted confocal microscope. Images were analyzed using ImageJ.

### Prolactin measurement

Conditioned-medium samples were collected on days 3, 5, and 10, stored at −80 °C, and analyzed using a Human Prolactin DuoSet ELISA kit (R&D Systems, catalog no. DY682). Capture antibody diluted in PBS was added to 96-well plates at 100 µL per well and incubated overnight at room temperature. Plates were washed three times with 400 µL of wash buffer per well and blocked with 300 µL of reagent diluent per well for at least 1 h.

Samples were diluted tenfold by combining 100 µL of sample with 900 µL of diluent. Diluted samples or standards were added at 100 µL per well and incubated for 2 h. Detection antibody was then added at 100 µL per well and incubated for 2 h, followed by 100 µL of streptavidin–horseradish peroxidase for 20 min, with aspiration and washing between steps. Substrate solution was added at 100 µL per well and incubated for 20 min protected from light. The reaction was stopped with 50 µL of stop solution per well. Absorbance was measured at 450 nm with wavelength correction at 540 or 570 nm, following the kit protocol. Prolactin concentrations were determined from the standard curve.

Comparisons used culture-area normalization based on 1.9 cm² for the two-dimensional 24-well culture and 1.09 cm² for the stromal-containing chip compartment. This approach assumed equal cell density at confluence. For comparisons among culture configurations, fold changes were calculated relative to the tri-culture prolactin value.

### Statistical analysis

Permeability and prolactin experiments were reported to include three biological replicates and three technical replicates. All other experiments were performed in technical triplicate. Permeability measurements were obtained from the same chips on days 5 and 10.

Data were summarized as mean ± standard deviation. The reported analysis used unpaired, two-tailed Student’s t-tests for two-group comparisons and one- or two-way analysis of variance followed by Tukey post hoc comparisons for multiple-group analyses. Statistical significance was defined as p < 0.05.

## Acknowledgements

Our work is supported by the Fonds de Recherche du Québec Santé (FRQS) (Chercheurs-boursiers J1 (313837) and Chercheurs-boursiers J2)), Establishment of Young Investigators (324277), Montreal TransMedTech Institute (iTMT), Start-up funds from the CRCHU Sainte-Justine (CRCHUSJ), University of Montreal, and Polytechnique Montreal to H.S

## Conflict of Interest

The Authors declare no conflict of interest.

## Author Contributions

**Ali Hedayatnia:** conceptualization, methodology, investigation, formal Analysis, visualization, validation, writing – original draft, writing – review and editing, data curation. **Arman Jafari:** methodology, formal Analysis, visualization, validation, writing – review and editing. **Nour Deca:** methodology, investigation, visualization. **Ramin Pajoomshariati:** co-supervision, conceptualization, methodology, investigation, formal Analysis, visualization, validation, writing – review and editing. **Houman Savoji:** conceptualization, methodology, validation, Supervision, funding acquisition, validation, project administration, resources, writing – review and editing.

## Data Availability

The data supporting the findings of this study are available within the article and upon request from the corresponding author.

## Table of Contents Text

VivoFemme combines a printed microfluidic device, a thin gelatin membrane, and gravity-driven flow to culture primary endometrial epithelial, stromal, and endothelial cells. The platform supports hormone-response and barrier-transport measurements across different culture configurations. Its integrated interface design provides a practical basis for investigating endometrial cell interactions, with further validation needed for predictive pharmacological applications.

## Notes

### Competing Interest Statement

The authors have declared no competing interest.

